# Demystifying the role of prognostic biomarkers in breast cancer through integrated transcriptome and pathway enrichment analyses

**DOI:** 10.1101/2022.06.20.496785

**Authors:** Divya Mishra, Ashish Mishra, M.P. Singh

## Abstract

Breast cancer (BC) is the most commonly diagnosed cancer and the leading cause of death in women. There has been discovered an increasing number of molecular targets for BC prognosis and therapy. However, it is still urgent to identify new biomarkers. Therefore, we evaluated biomarkers that may contribute to the diagnosis and treatment of BC. We searched TCGA datasets and identified differentially expressed genes (DEGs) by comparing tumor (100 samples) and non-tumor (100 samples) tissues using the Deseq2 package. Pathway and functional enrichment analysis of the DEGs were done using *DAVID (*The *Database* for Annotation, Visualization, and Integrated Discovery*) database*. The protein-protein interaction (PPI) network was identified using the STRING database and visualized through Cytoscape software. Hub gene analysis of the PPI network was done using Cytohubba plugins. The associations between the identified genes and overall survival (OS) were analyzed using Kaplan–Meier plot. Finally, we have identified hub genes at the transcriptome level. A total of 824 DEGs were identified, which were mostly enriched in cell proliferation, signal transduction, and cell division. The PPI network comprised 822 nodes and 12145 edges. Elevated expression of the 5 hub genes AURKA, BUB1B, CCNA2, CCNB2, and PBK are related to poor OS in breast cancer patients. A promoter methylation study showed these genes to be hypomethylated. Validation through genetic alteration and missense mutations resulted in chromosomal instability leading to improper chromosome segregation causing aneuploidy. The enriched functions and pathways included the cell cycle, oocyte meiosis, and the p53 signaling pathway. The identified five hub genes in breast cancer have the potential to become useful targets for the diagnosis and treatment of breast cancer.

## Introduction

Breast cancer (BC) is the most common type of cancer and the second most prominent cause of cancer-related death in women (Zeng et al. 2021). According to the World health organization (WHO), in 2020, there were 2.3 million women diagnosed with breast cancer and 685000 deaths globally (WHO, 2020). The lack of improved adjuvant therapy is also a major problem in reducing the burden of BC patients. Currently, the lymph node involvement, tumor size, and distant metastasis of the American Joint Committee on Cancer have been extensively identified, but there is still a need for a globally recognized platform or efficient markers that can correctly predict the prognosis of BC patients (Pan et al. 2017). Even though applying for endocrine therapy or neoadjuvant chemotherapy, clinic-pathological parameters are commonly ambiguous, which complicates the judgments of real prognosis (Deng et al. 2019). Approximately 70-80% of BC patients can be cured, especially when the disease identified early, while advanced BC having distant organ metastases is considered incurable with currently available treatment strategies. Therefore, there is a critical need to find breast cancer biomarkers that can help to develop better treatment strategies for breast cancer. Comprehensive research required to focus on understanding the molecular basis of BC (Kim et al. 2020).

Since then, many genes have been identified as prognostic and predictive biomarkers of breast cancer that play a significant role in precise treatment (Ellsworth et al. 2011; Zeng et al. 2021). The commonly targeted drugs used for HER2-positive BC include trastuzumab, lapatinib, tucatinib, trastuzumab emtansine (T-DM1) and pertuzumab. Many molecular-targeted drugs therapy include mammalian target of rapamycin (mTOR)/ serine/threonine kinase (AKT)/ phosphoinositide 3-kinase (PI3K) signaling pathways, which include bupacoxib, abencoxib, GDC-0068, alpelisib, and Bez235 (Zeng et al. 2021). Therefore, vascular endothelial growth factor has found to be as a key target for anti-angiogenic treatement, and its reported inhibitors such as sorafenib, sunitinib, and bevacizumab are utilizing for breast cancer therapy (Zeng et al. 2021). Androgen receptor (AR) based targeted therapies, can include AR antagonists and AR agonists which showing prominent results in clinical trials for BC patients (Jin et al. 2019; Zeng et al. 2021).

Likewise, the combinations of AR-based targeted treatments with other reagents such as PI3K inhibitor have been analyzed to overcome resistance to AR-targeted treatments. In contrast, the targeted treatment strategies have been extensively developed for cyclin-dependent kinase 4/6 (CDK4/6), *BRCA1/2*-mutated polyadenosine diphosphate ribose polymerase (PARP), BTB and CNC homology1 (BACH1), epidermal growth factor receptor (EGFR) and so on. However, due to low ratios of responders, tumor heterogeneity, drug resistance, there is still a strong need to identify new biomarkers that can help to diagnose and treat BC (Zeng et al. 2021).

Computational analysis is the efficient strategies for the comprehensive study of large databases that include complex genomic information (Ren et al. 2020). Our present study used sophisticated *in silico* approaches to identify potential prognostic biomarkers that can be useful for BC. This analysis, therefore include the identification of differentially expressed genes that were overexpressed in BC. The 5 hub genes obtained were further validated through promoter methylation, mutation and genetic alterations analysis proved their potential to be prognostic biomarkers. The survival analysis of all these hub genes showed poorer survival rates among BC patients.

## Materials and Methods

### Fetching and preprocessing of data

The raw data for the solid normal samples and primary tumor was obtained from The Cancer Genome Atlas (TCGA). The raw data was pre-processed using bioinformatics tools and software. The quality assessment of the raw reads was carried out using FastQC (v 0.11.8) to identify the short length reads (adapter content) having low quality and uncalled biases. The low quality reads were filtered and trimmed using Cutadapt software tool (v 3.2) for removing the noise in the data that could affect the results drastically. The trimmed reads were further aligned against the human reference genome (GRch38/hg38) using STAR alignment tool (v 2.7.7a) and is considered as one of the fastest global alignment tool (Dobin et al., 2013). In the next step, the mapped reads were quantified to obtain the read counts corresponding to each gene through featureCounts (v 2.0.1) (Liao et al., 2013). Finally, the differentially expressed genes (DEGs) were obtained between solid normal samples and primary tumor through DESeq2 (v 1.22.1) which provided the quantitative variation in the expression levels of genes. This process is based on the normalization of the data using negative binomial distribution (Love et al., 2014). The criteria specified for categorizing the genes as significantly differentially expressed was false discovery rate (p-value (adj.) < 0.05) and |log_2_FC| > 2.

The flowchart shown below depicts the entire process that was followed in this study (Figure 1).

**Figure 1.**
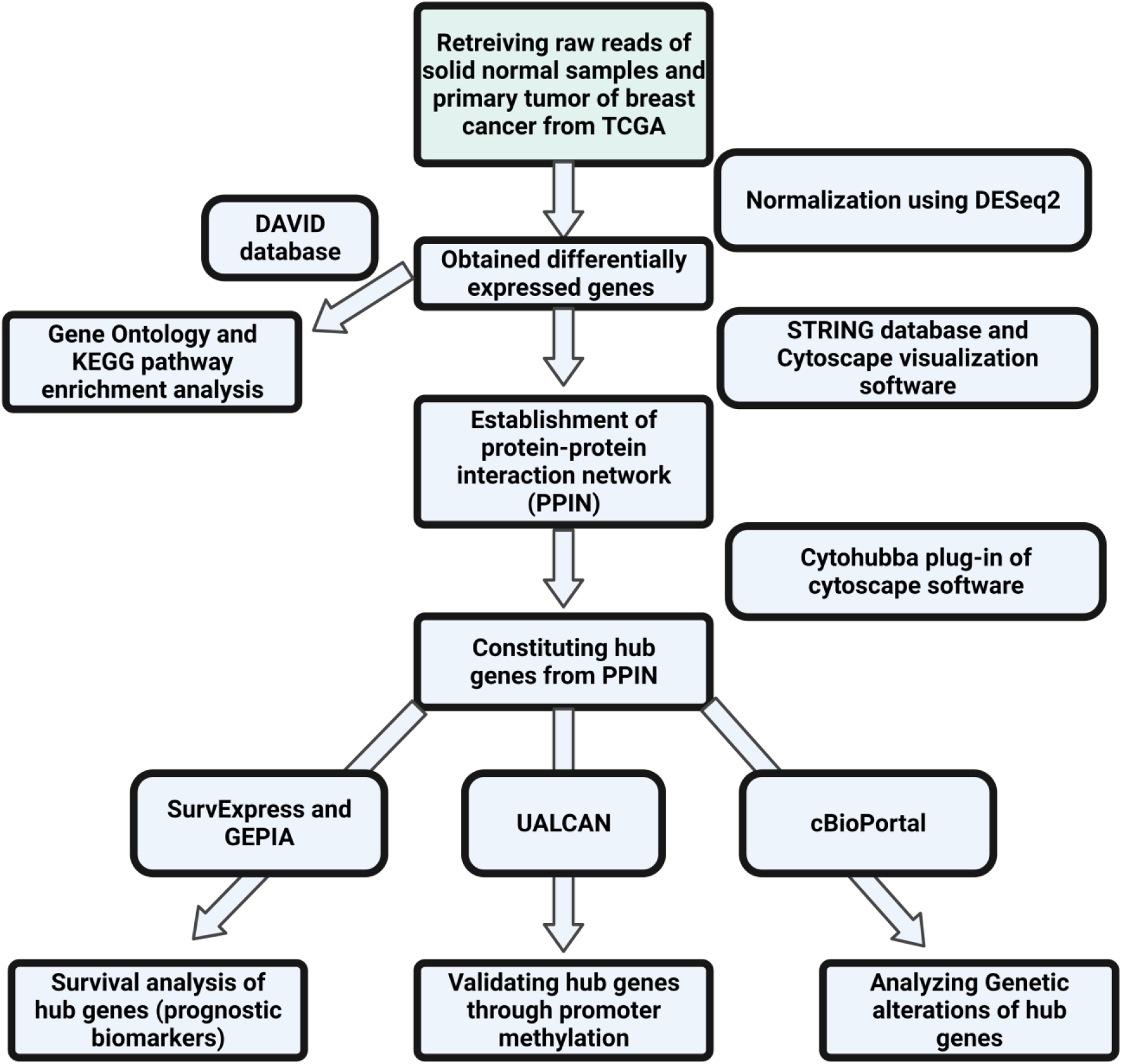
Flowchart showing the methodology followed in the present study.

### Investigating the protein-protein interaction network (PPIN) to establish the hub genes as potential prognostic biomarkers

The protein-protein interaction network deals with mathematical representations pertaining to physical contacts established between different cellular level proteins and are crucial to understand the processes that are taking place at the cellular level in normal and diseased state. STRING database developed for the purpose of constructing the PPI network was used in this case and this database uses the differentially expressed genes as input to provide the required result (Szclarczyk et al., 2020). The nodes of the network correspond to differentially expressed genes (DEGs) and the edges constitute the interaction between the proteins. Cytoscape visualization software was used to visualize the various interactions and analyzing the PPI network (Shannon, P. et al., 2003). The significance of the interactions in the PPI network was analyzed through PPI enrichment value < 1.0e-16. Confidence interval <0.4 was set for constructing the PPIN. For determining the hub genes as prognostic biomarkers, cytohubba plug-in, available in the cytoscape software, was used. 6 significant topologies of cytohubba viz. Degree, Maximal Clique Centrality (MCC), Maximum Neighborhood Component (MNC), Edge Percolated Component (EPC), Radiality, and closeness were employed. From these five algorithms, the hub genes common among all of these were finally established using jVenn online tool (Bardou et al., 2014).

### Analyzing the gene ontology (GO) components and enriched pathways involved in the progression of breast cancer

DAVID (Database for Annotation, Visualization and Integrated Discovery) is an online tool for establishing the functional enrichment of overexpressed genes involved in different disease types (Dennis Jr et al., 2003). In case of the present study, the gene list was uploaded in the database for exploring both GO terms and KEGG pathways involved in breast cancer. The modified Fisher Exact P-value was set to 0.1 and this value aid in the measurement of gene-enrichment in annotation terms. Likewise, the value for count threshold was fixed at 2 and this is the default value in the database. The lesser value of p-value is indicates more enriched GO terms and KEGG pathways. These terms are considered significant based on the cut-off value for any term or pathway which was set at p < 0.05. For visualizing these obtained components from DAVID, an online server, REVIGO (Supek et al., 2011) was used. It provided the treemaps corresponding to biological processes, cellular components and molecular functions based on the GO ids and respective p-values of each component.

### Exploring the epigenetic regulation of hub genes through promoter methylation

The analysis of the consequences on the overexpressed genes due to the variations at the epigenetic level provides an in-depth knowledge about tumorigenesis and metastasis of breast cancer. Promoter methylation study provides this information and it can be obtained for each gene through an online server, UALCAN (Chandrashekar et al., 2017). This multi-omics server dedicated to cancer study employs TCGA datasets and for the analysis of the present study, datasets related to breast cancer was employed. The result could be interpreted based on the beta values that indicate the level of DNA methylation. These values range from 0 (unmethylated) to 1 (fully-methylated). The beta values having range between 0.5 and 0.7 pertains to hypermethylation, while those between 0.05 and 0.3 corresponds hypomethylation.

### Identifying the genetic alterations of hub genes

Different external and internal factors are responsible for causing genetic alterations such as mutations and copy number alterations and these alterations results in altering the DNA sequences and plays a pivotal role in the development and progression of cancer, its metastasis and providing resistance to therapies. In the present study, these genetic alterations in the hub genes were identified using cBioPortal online resource which contains genomic datasets of patients suffering from different cancer types (Gao et al., 2013). The results pertaining to copy number alterations were obtained from GISTIC (Genomic Identification of Significant Targets in Cancer) algorithms which identifies the significantly altered regions across the different sets of patients. These results obtained from GISTIC correspond to the level of copy-number per gene where a value of -2 indicates deep deletion or deep loss and constitutes homozygous deletion. Similarly, a value of -1 corresponds to shallow deletion and constitutes a heterozygous deletion. The value 0 attributes to normal or diploid, 1 to gain (low-level gain) and 2 to amplification (high-level amplification). For visualizing these alterations (mutations and copy number alterations) obtained for different hub genes, OncoPrints was used. The mutations occurred in the intronic region referred to splice site mutation while that occurred at the exon/intron junction referred to splice region mutations.

### Validating the differential expression pattern and survival analysis of hub genes

GEPIA (Gene Expression Profiling Interactive Analysis), an online web server (Tang et al., 2017) was used to obtain the gene expression profiles of all the 5 hub genes in case of patients suffering from breast cancer. The survival analysis corresponding to these hub genes were obtained from SurvExpress (Aguirre-Gamboa et al., 2013). The Kaplan-Meier (KM) plot used for visualizing the survival analyses of all the hub genes (prognostic biomarkers) is based on the univariate Cox regression analysis which provides the risk score by categorizing the patients into low- and high-risk groups.

## Results

### Determination of differentially expressed genes through statistical analysis

The RNA-Seq high-throughput analysis produced 2854 differentially expressed genes (DEGs) for breast cancer out of which 1812 were upregulated and 1042 were downregulated. The upregulated and downregulated genes can be visualized using Bland-Altman (MA) plot (Figure 2(a)). It could be evidenced from the figure that more number of DEGs was found in the positive x-axis showing more upregulated genes as compared to the downregulated genes in the negative x-axis. The volcano plot (Figure 2(b)) that provides the information about the most significant differentially expressed genes showed that all the 5 identified biomarkers in this study were upregulated as they all lie on the left portion of the plot and shown by red dots. The most significant differentially expressed gene among these 5 DEGs was BUB1B having the highest log fold change value in the deseq2 statistical analysis.

**Figure 2.**
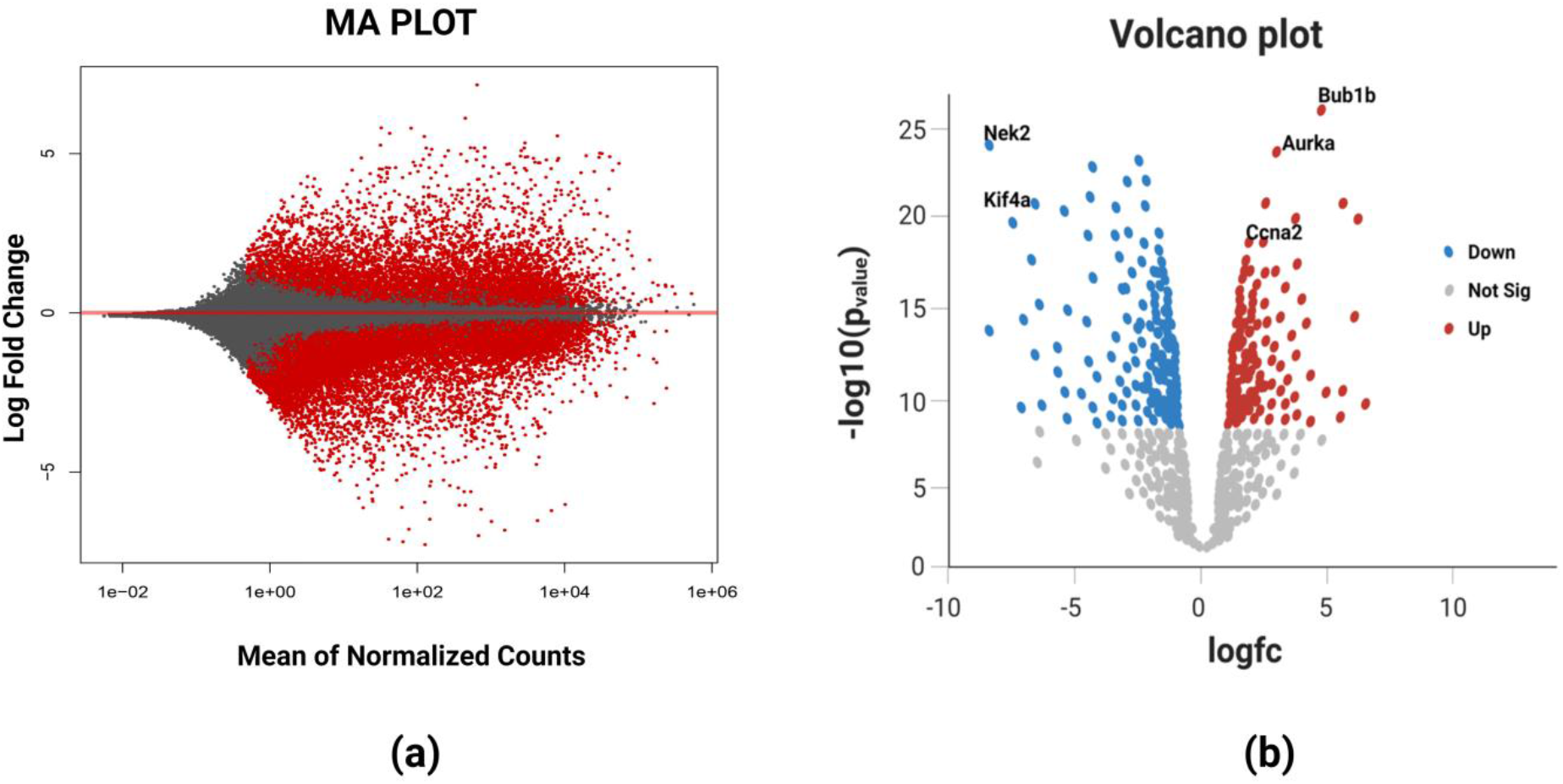
(a) MA plot of breast cancer representing the log fold-change against mean expression using DESeq2 dataset. The red dots corresponding to positive x-axis represents upregulated differentially expressed genes while those corresponding to negative x-axis represent downregulated differentially expressed genes. (b) Volcano plot showing the most significant differentially expressed genes. The blue color dots on the left portion of the plot represents the downregulated DEGs, red color dots on the right portion of the plot represents upregulated DEGs and white color dots at the bottom portion depicts the non-significant DEGs. BUB1B is the most significant DEG based on its highest value of log fold change.

### Investigation of the protein-protein interaction network (PPIN) established the hub genes as potential prognostic biomarkers

The obtained DEGS were used for constructing the PPIN having 822 nodes, and 12145 edges. The average node degree was 29.5, average local clustering coefficient was 0.453, and PPI enrichment p-value was less than 1.0e-16. The PPIN with the above characteristics is shown below (Figure 3). The 5 hub genes obtained from different topologies of cytohubba are-AURKA (Aurora Kinase A), BUB1B (BUB1 Mitotic Checkpoint Serine/Threonine Kinase B), CCNA2 (Cyclin A2), CCNB2 (Cyclin B2), and PBK (PDZ Binding Kinase) (Figure 4). The values and ranks of the hub genes in these algorithms are summarized in the table (Table 2). The 5 hub genes were upregulated in breast cancer and promoting tumorigenesis and metastasis.

**Table 2.** Table showing the values of hub genes for various topological algorithms of cytohubba.

| Hub Genes | Closeness | Degree | EPC | MCC | MNC | Radiality |
| --- | --- | --- | --- | --- | --- | --- |
| AURKA | 192.6 | 106 | 38.26 | 1.65e+57 | 106 | 6.67 |
| BUB1B | 194.7 | 112 | 43.76 | 1.65e+57 | 112 | 6.62 |
| CCNA2 | 200.4 | 109 | 38.26 | 1.65e+57 | 109 | 6.73 |
| CCNB2 | 190.6 | 110 | 41.25 | 1.65e+57 | 110 | 6.52 |
| PBK | 199.5 | 115 | 42.23 | 1.65e+57 | 115 | 6.61 |

**Figure 3.**
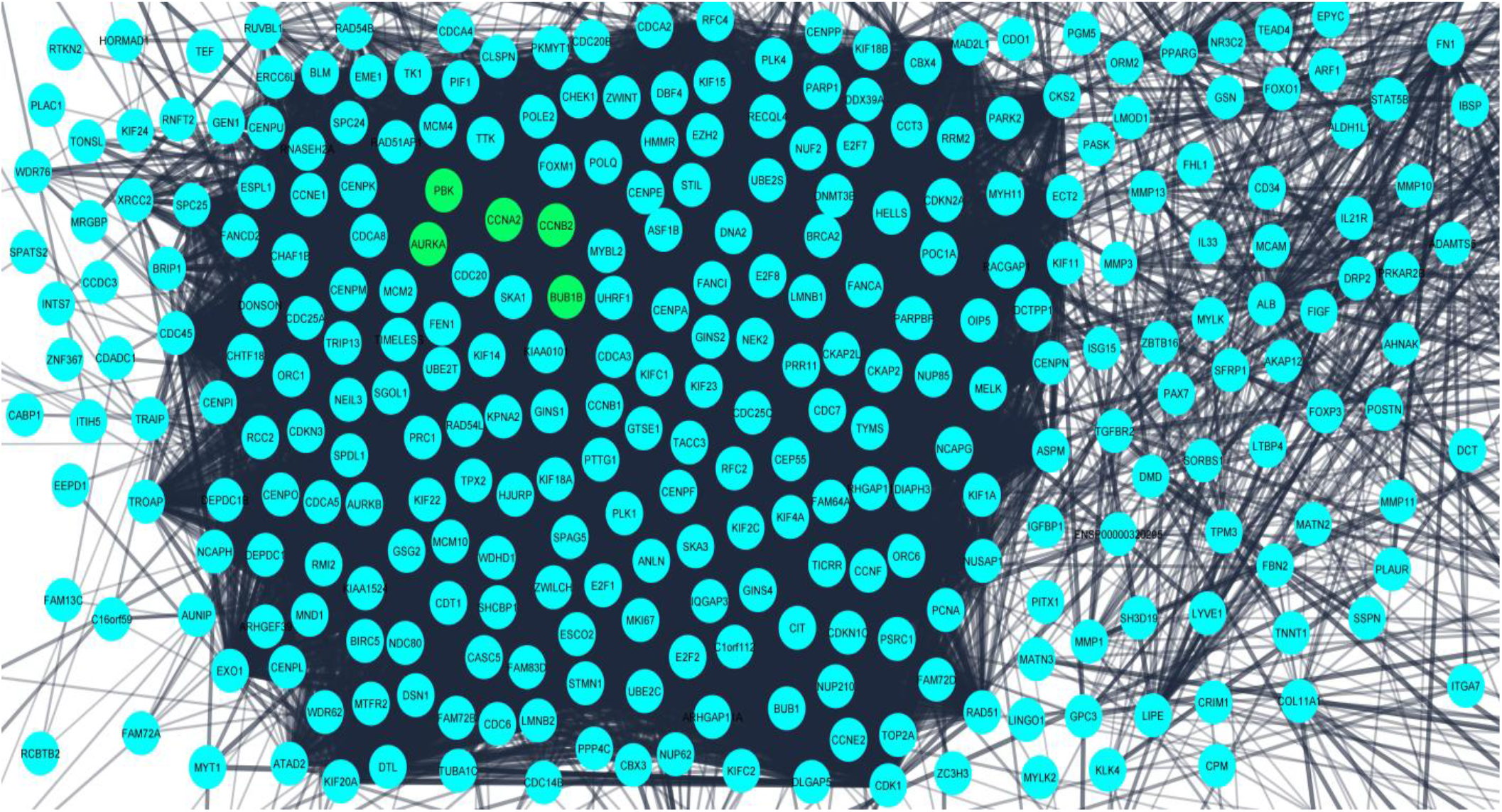
Protein-Protein Interaction Network for the 824 differentially expressed genes obtained from STRING database. The confidence interval was > 0.7. Nodes represent interacting proteins and the edges represent the interaction between these proteins. The 5 hub genes (potential prognostic biomarkers) are shown in green color.

**Figure 4.**
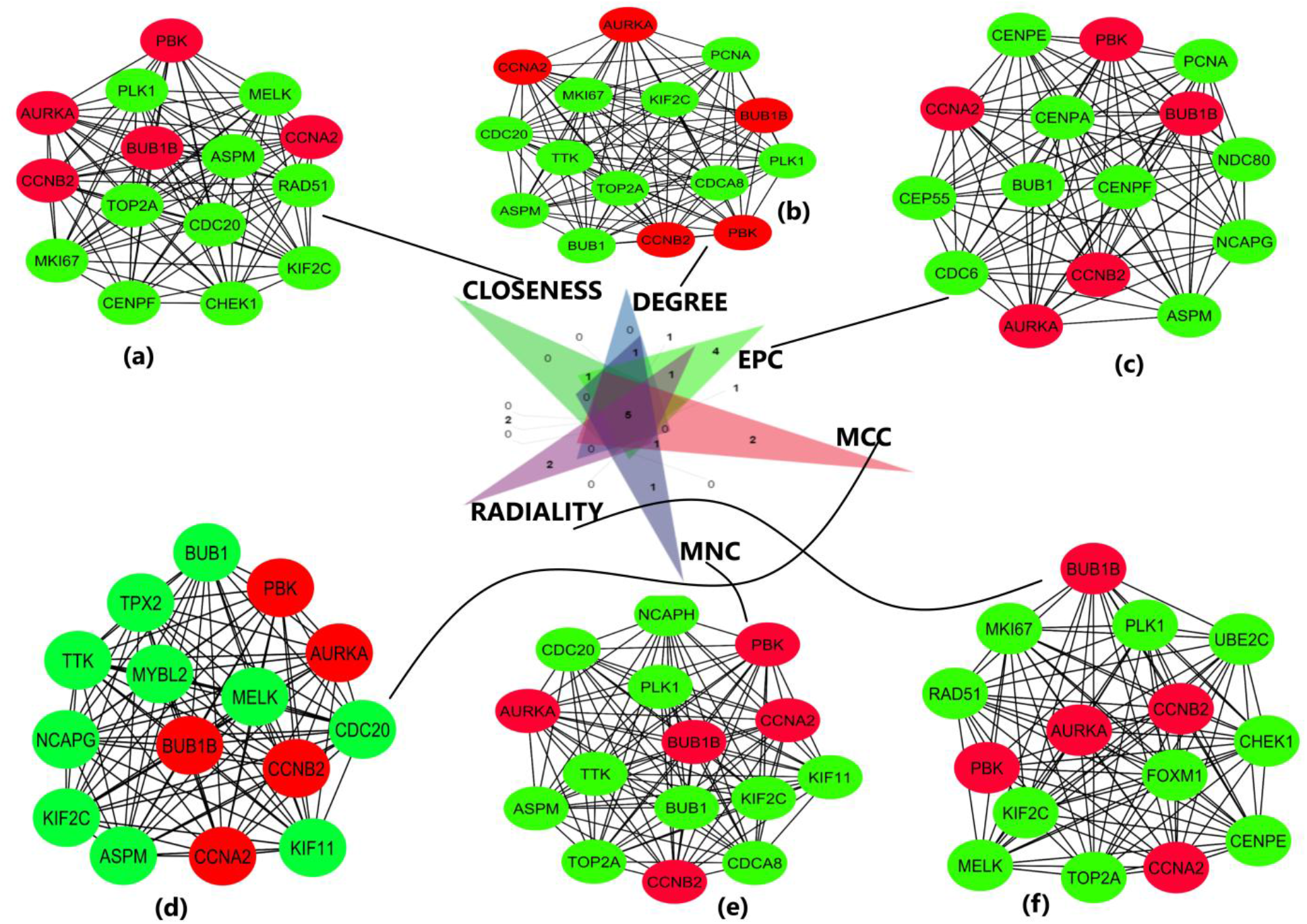
Important Sub-Networks and nodes obtained from cytohubba plug-in of cytoscape software using six topological algorithms. The Top 15 hub genes were evaluated in the PPI network using these six calculation methods. (a) Sub-network obtained from closeness topological algorithm. The nodes in red represent the top-ranked hub genes (b) Sub-network and hub genes obtained from degree topological algorithm (c) Sub-network and hub genes obtained from EPC topological algorithm (d) Sub-network and hub genes obtained from MCC topological algorithm (e) Sub-network and hub genes obtained from MNC topological algorithm (f) Sub-network and hub genes obtained from radiality topological algorithm. The hub genes are shown in red color in all the 6 topologies.

### Gene Oncology (GO) Component and KEGG Pathway Enrichment Analysis

The DAVID database provided the components and pathways in which the 5 hub genes were participated and enriched. The hub genes were found to be enriched in various biological processes as cell cycle, mitotic cell cycle, cell division, mitotic nuclear division, and chromosome segregation and these are some of the most important processes that promotes tumorigenesis and metastasis of breast cancer (Figure 5). The biological processes were ranked based on p-values and these processes along-with their respective p-values are tabulated in the table (Table 3).

**Table 3.** Top 10 significantly enriched biological processes along-with their respective p-values.

| Biological Process | p-value |
| --- | --- |
| GO:0000278~Mitotic Cell Cycle | 1.13e-43 |
| GO:1903047~Mitotic Cell Cycle Process | 2.43e-43 |
| GO:0022402~Cell Cycle Process | 7.17e-40 |
| GO:0007049~Cell Cycle | 1.97e-35 |
| GO:0000280~Nuclear Division | 1.99e-33 |
| GO:0051301~Cell Division | 3.12e-33 |
| GO:0007059~Chromosome Segregation | 4.28e-33 |
| GO:0000819~Sister Chromatid Segregation | 5.08e-33 |
| GO:0007067~Mitotic Nuclear Division | 3.20e-32 |
| GO:0048285~Organelle Fission | 1.05e-31 |

**Figure 5.**
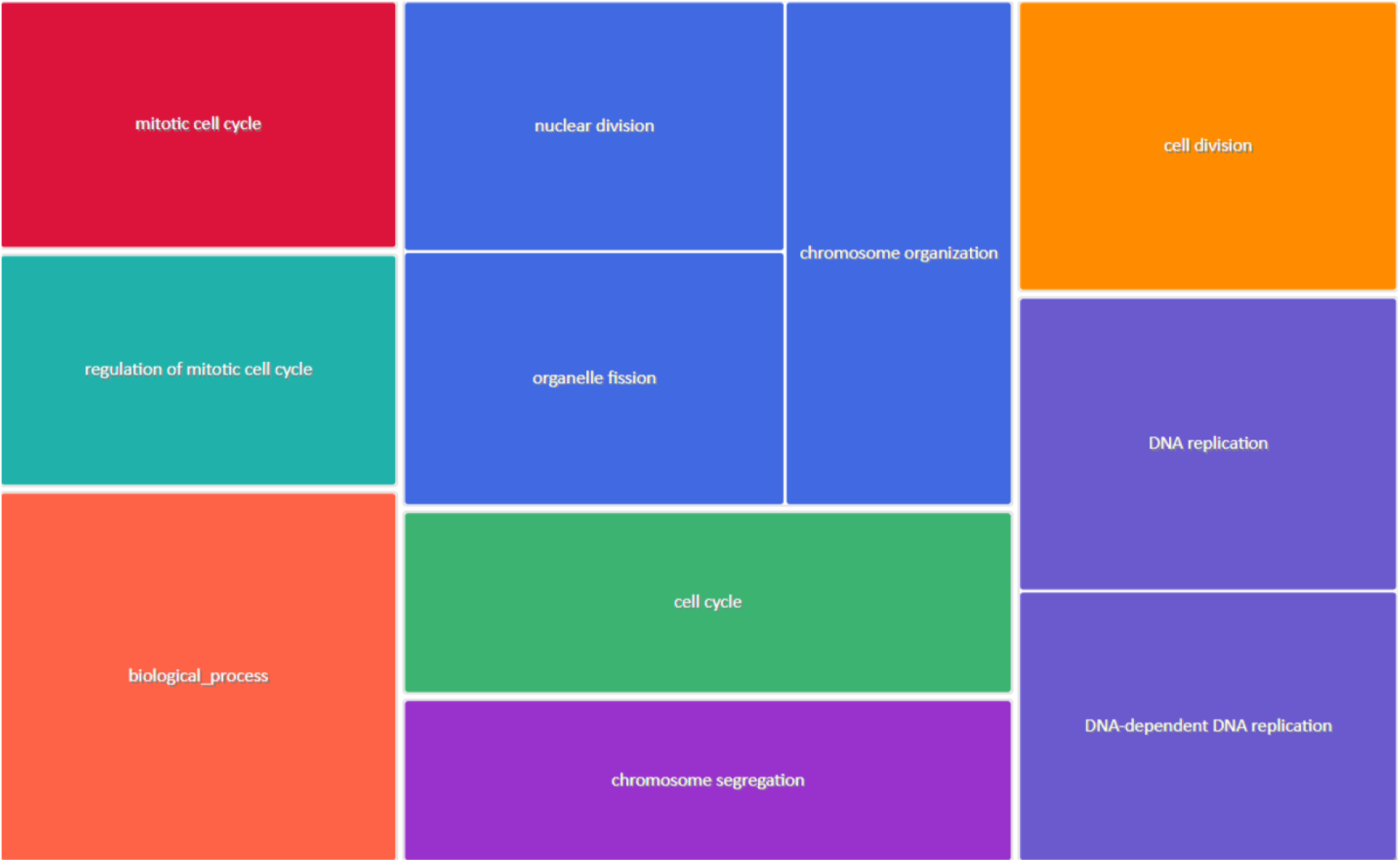
Biological Processes (BP) based on p-values drawn from Revigo in which the hub genes were significantly enriched.

The significant KEGG pathways based on p-values include oocyte meiosis, cell cycle, progesterone-mediated oocyte maturation, and p53 signaling pathway (Figure 6). Some of the top-ranked enriched KEGG pathways along with their respective p-values are tabulated below (Table 4).

**Table 4.**
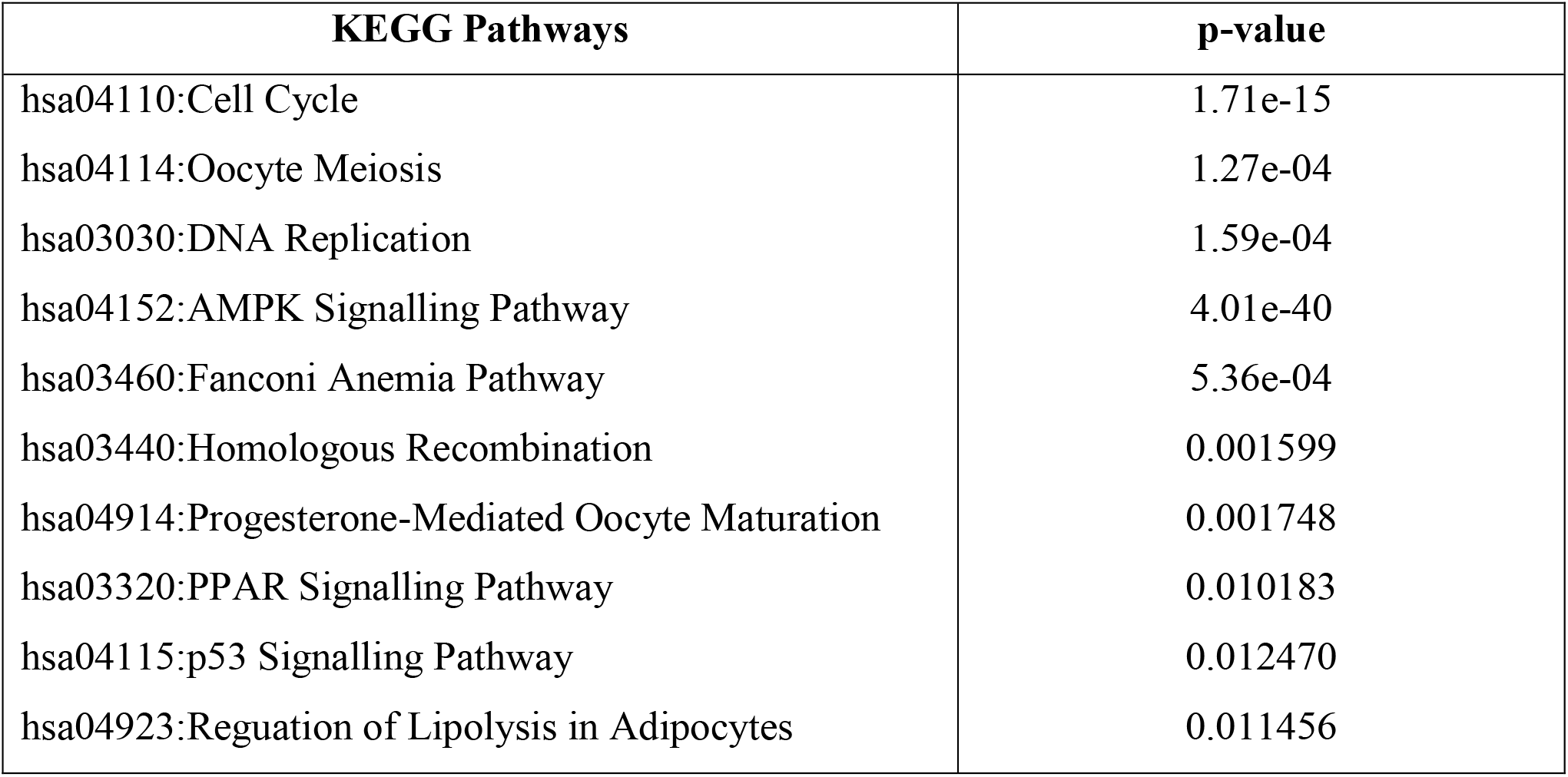
Top 10 significantly enriched KEGG pathways with their respective p-values.

| **KEGG Pathways** | **p-value** |
| --- | --- |
| hsa04110:Cell Cycle | 1.71e-15 |
| hsa04114:Oocyte Meiosis | 1.27e-04 |
| hsa03030:DNA Replication | 1.59e-04 |
| hsa04152:AMPK Signalling Pathway | 4.01e-40 |
| hsa03460:Fanconi Anemia Pathway | 5.36e-04 |
| hsa03440:Homologous Recombination | 0.001599 |
| hsa04914:Progesterone-Mediated Oocyte Maturation | 0.001748 |
| hsa03320:PPAR Signalling Pathway | 0.010183 |
| hsa04115:p53 Signalling Pathway | 0.012470 |
| hsa04923:Regulation of Lipolysis in Adipocytes | 0.011456 |

**Figure 6.**
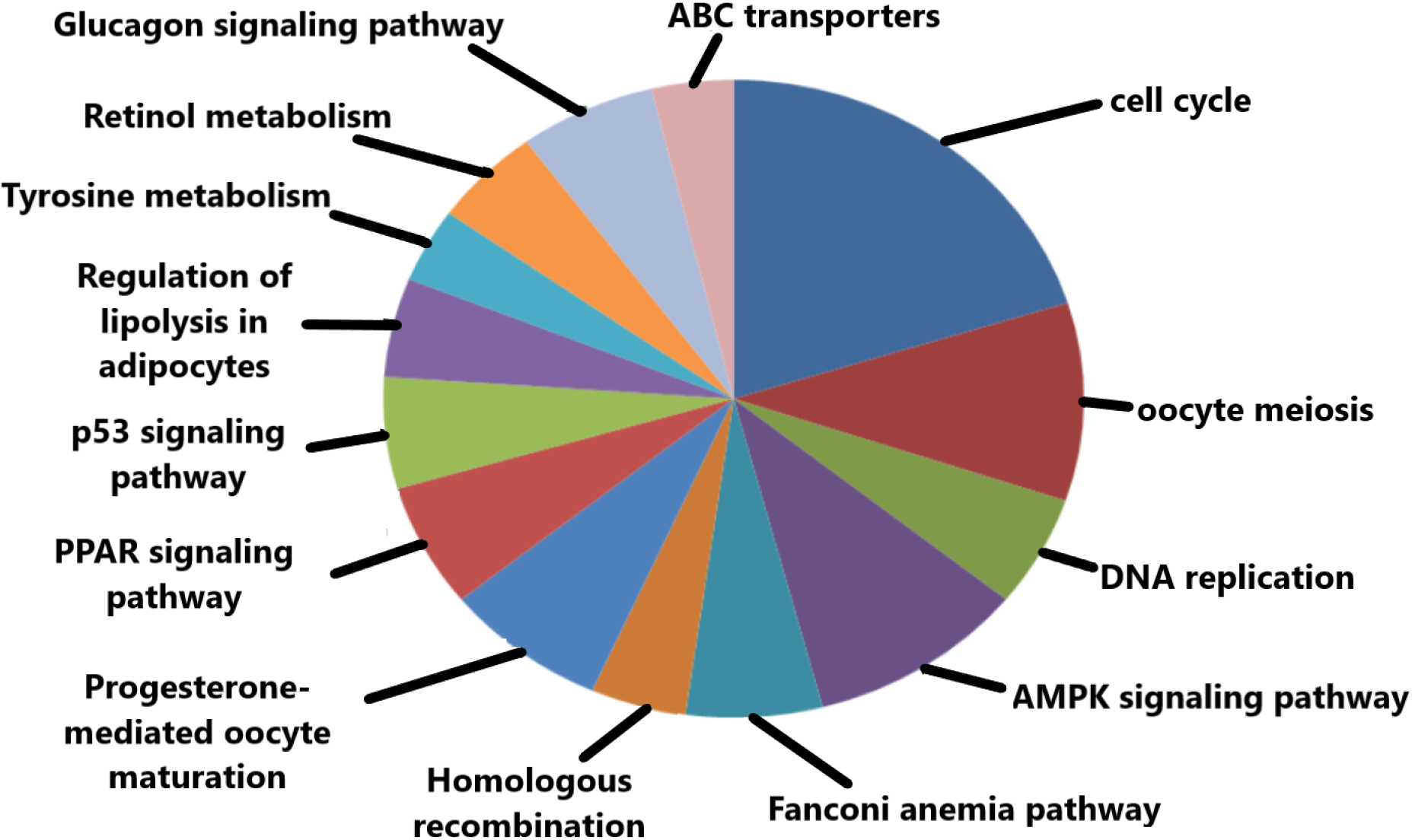
Important KEGG pathways in which the hub genes were significantly enriched.

### Exploring the epigenetic regulation of hub genes through promoter methylation

Validation of promoter methylation using UALCAN database revealed that the promoter methylation level of BUB1B and CCNB2 was lower than normal samples in breast cancer that indicates higher expression of these hub genes (Figure 7 (b) & (d)) in contrast to that of AURKA, CCNA2 and PBK having higher promoter methylation level than normal samples (Figure 7 (a), (c) & (e)).

**Figure 7.**
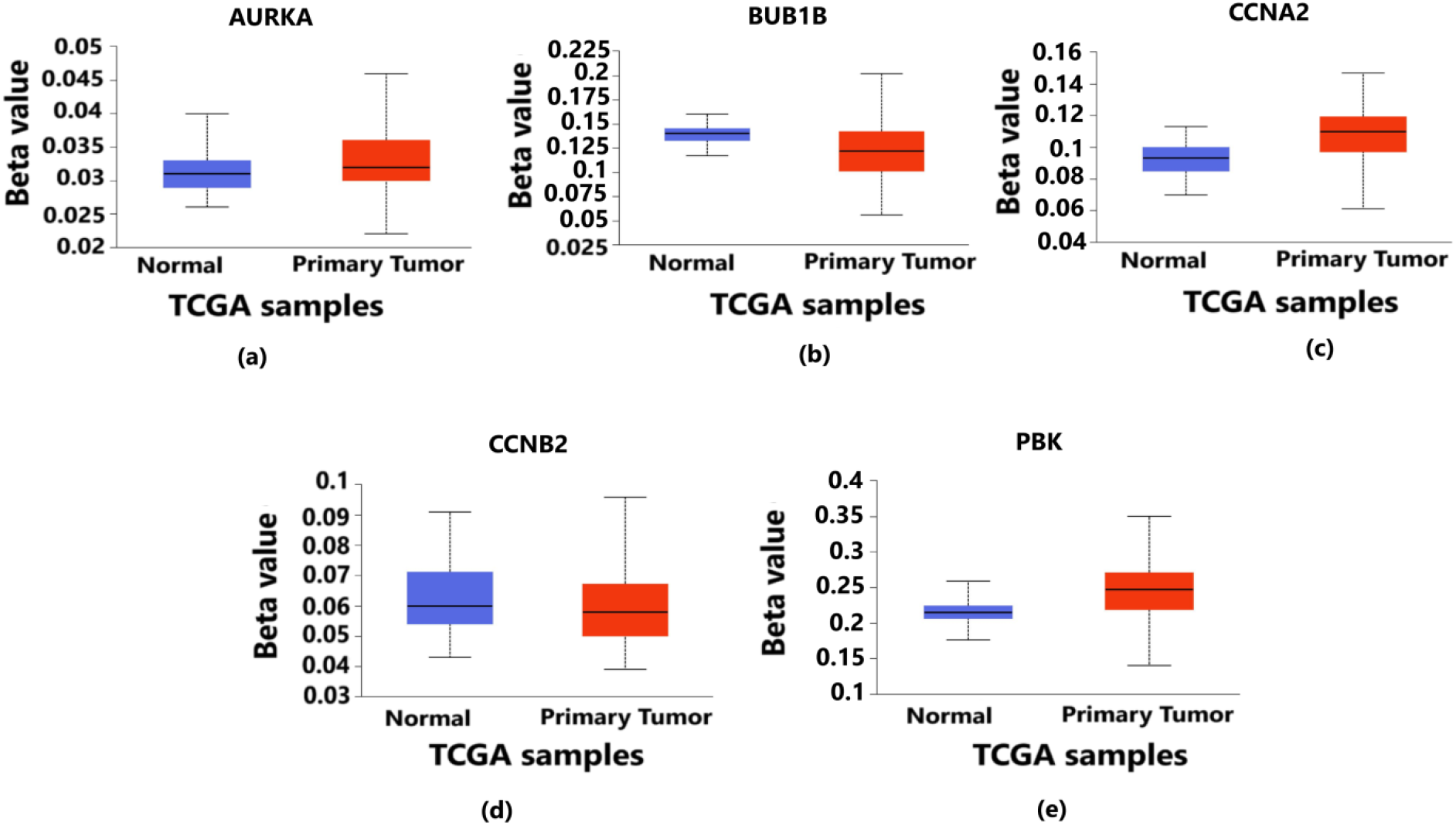
Level of Promoter methylation corresponding to 5 hub genes in Breast Cancer. It provides the information about whether the over-expressed genes are hypermethylated or hypomethylated as both plays vital role in the progression and metastasis of breast cancer. (a) Methylation level of AURKA gene in which the methylation level of tumor samples are higher than normal samples (b) Methylation level of BUB1B gene in which the methylation level of tumor samples are lower than normal samples (c) Methylation level of CCNA2 gene in which the methylation level of tumor samples are higher than normal samples (d) Methylation level of CCNB2 gene in which the methylation level of tumor samples are lower than normal samples (e) Methylation level of PBK gene in which the methylation level of tumor samples are higher than normal samples

### Findings of genetic alterations in hub genes

Tumorigenesis mainly occurs due to irremediable mutations in cell structures. These mutations could be identified through genetic alteration analysis. The alterations may be in the form of missense mutation, splice mutation, deep deletion, truncating mutation, and amplification. In case of breast cancer, the percentage alteration of all the 5 hub genes varied from 0.7% to 6% (Figure 8 (a)). The corresponding frequency of occurrence of the genetic alterations shows more frequency of amplification and mutations in all the 5 hub genes (Figure 8 (b)). Copy number alterations for breast cancer show most of the alterations due to diploid, gain, and amplification. AURKA gene was mostly affected due to amplification in the genetic materials while the remaining 4 hub genes were mainly altered due to either gain, diploid or in some cases, deep deletion (Figure 9). The details of genetic alterations and copy number variations are summarized in the table below (Table 5). Almost all the mutations in these 5 hub genes consist of phosphorylation PTMs.

**Table 5.**
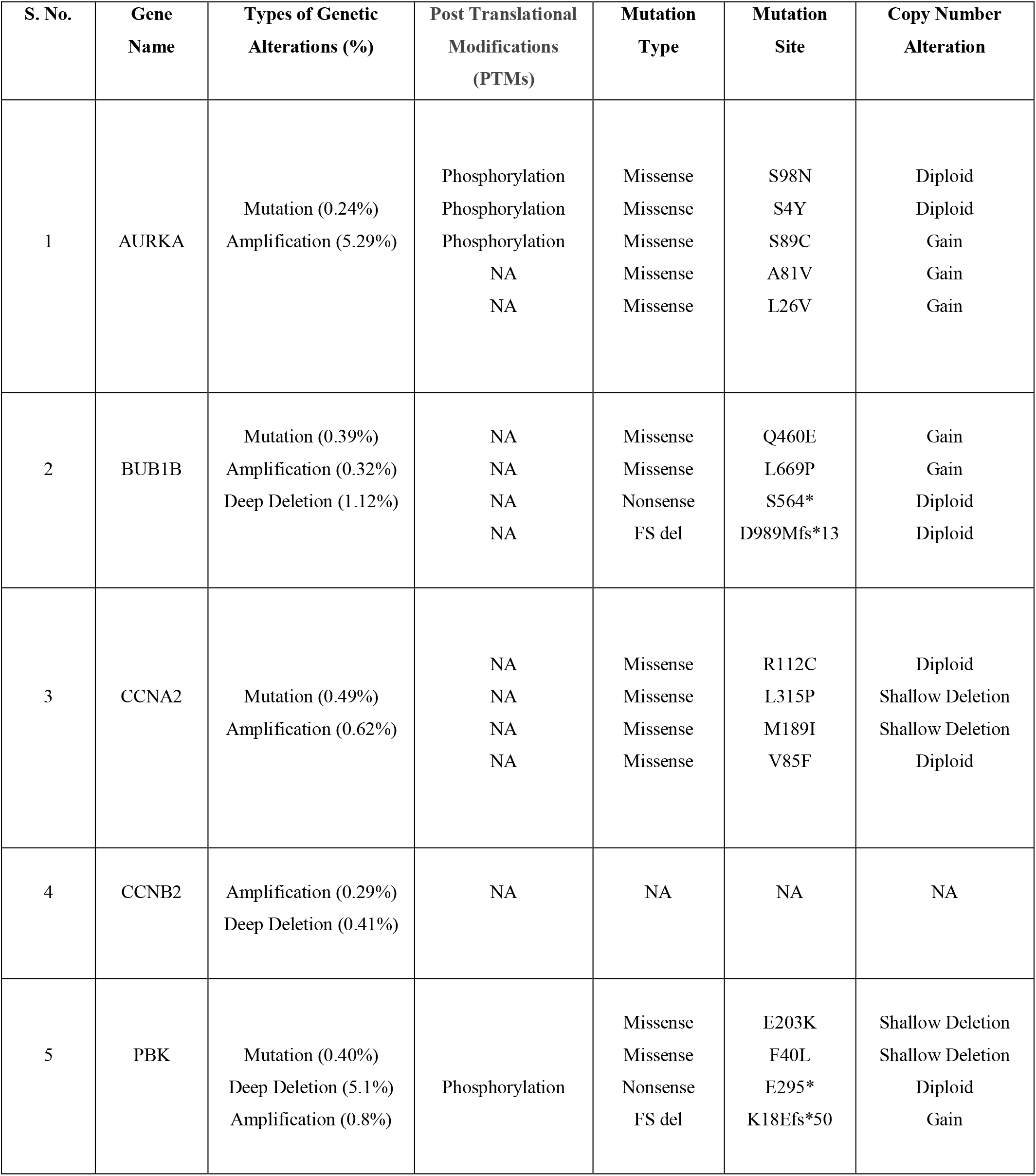
Table summarizing the information related to genetic alterations in breast cancer.

**Figure 8.**
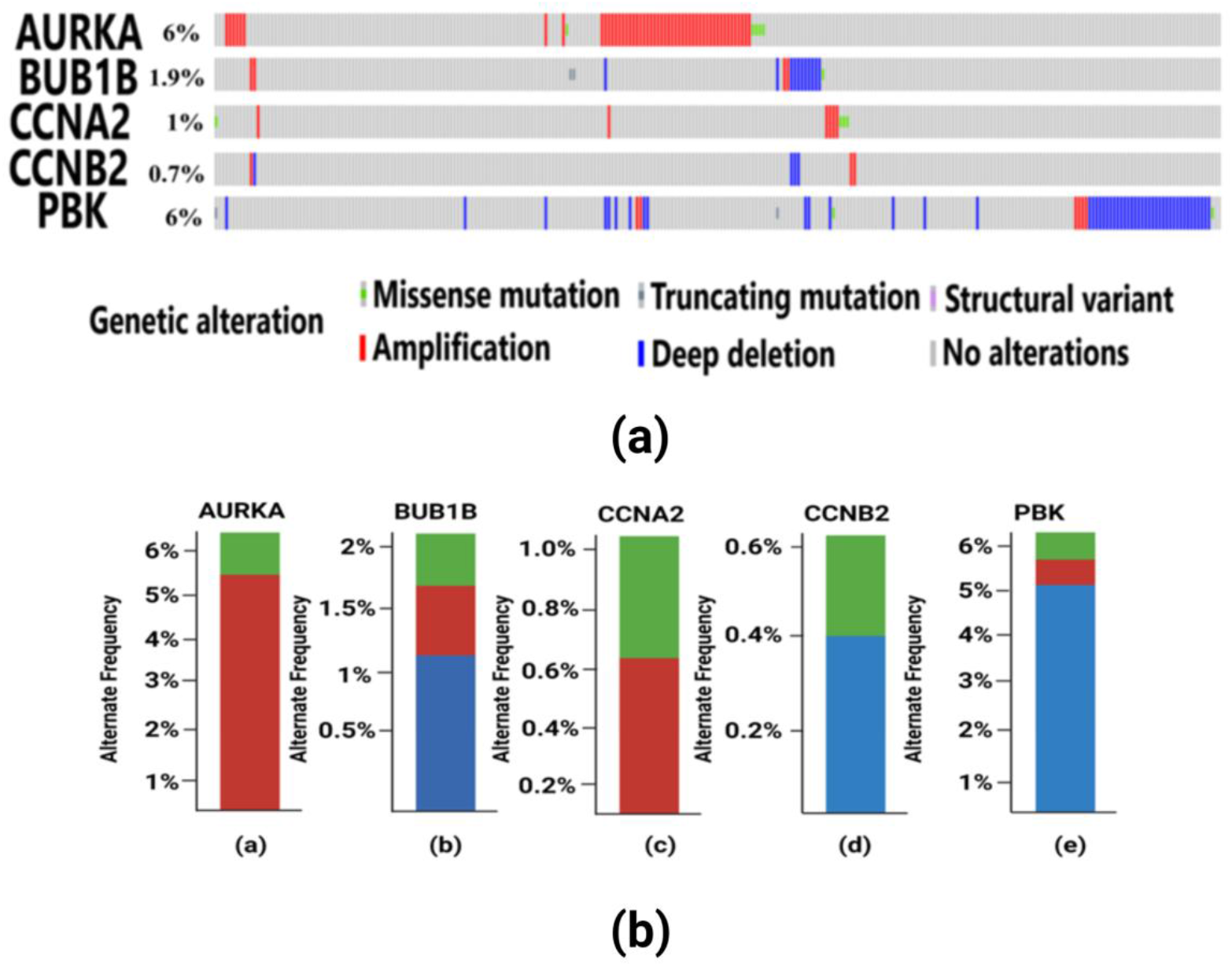
(a) Visualization of genetic alterations of hub genes in breast cancer using OncoPrint. In this figure, AURKA have 6% genetic alteration having missense mutation in 5 TCGA patient samples and amplification in other samples. BUB1B have 1.9% genetic alterations which include missense and truncating mutations, amplification and deep deletion in some samples. CCNA2 have 1% genetic alterations having missense and amplification. CCNB2 consists of 0.7% alterations mainly consists of amplification and deep deletion. PBK gene have 6% alterations having truncating and missense mutations, amplification and deep deletion in the patient samples. (b) Frequency of genetic alterations in hub genes in breast cancer. Red color indicates amplification, green color indicates mutations and blue color indicates deep deletion. (a) AURKA gene has more frequent occurrence of amplification in 5% of the samples and less frequent mutations in only 1% of the samples (b) BUB1B gene has more frequent deep deletion in 1% of the samples, followed by less frequent amplification and mutation each occurring in only 0.5% of the samples (c) CCNA2 gene has more frequency of amplification in 0.6% of the samples and mutations in 0.4% of the samples (d) CCNB2 gene has deep deletion having frequent occurrence in 0.4% of the samples and less frequency of occurrence of amplification in 0.2% of the patient samples (e) PBK gene has more genetic alterations due to deep deletion in 5% of the samples, followed by amplification in 0.8% of the samples and mutation in 0.2% of the samples respectively.

**Figure 9.**
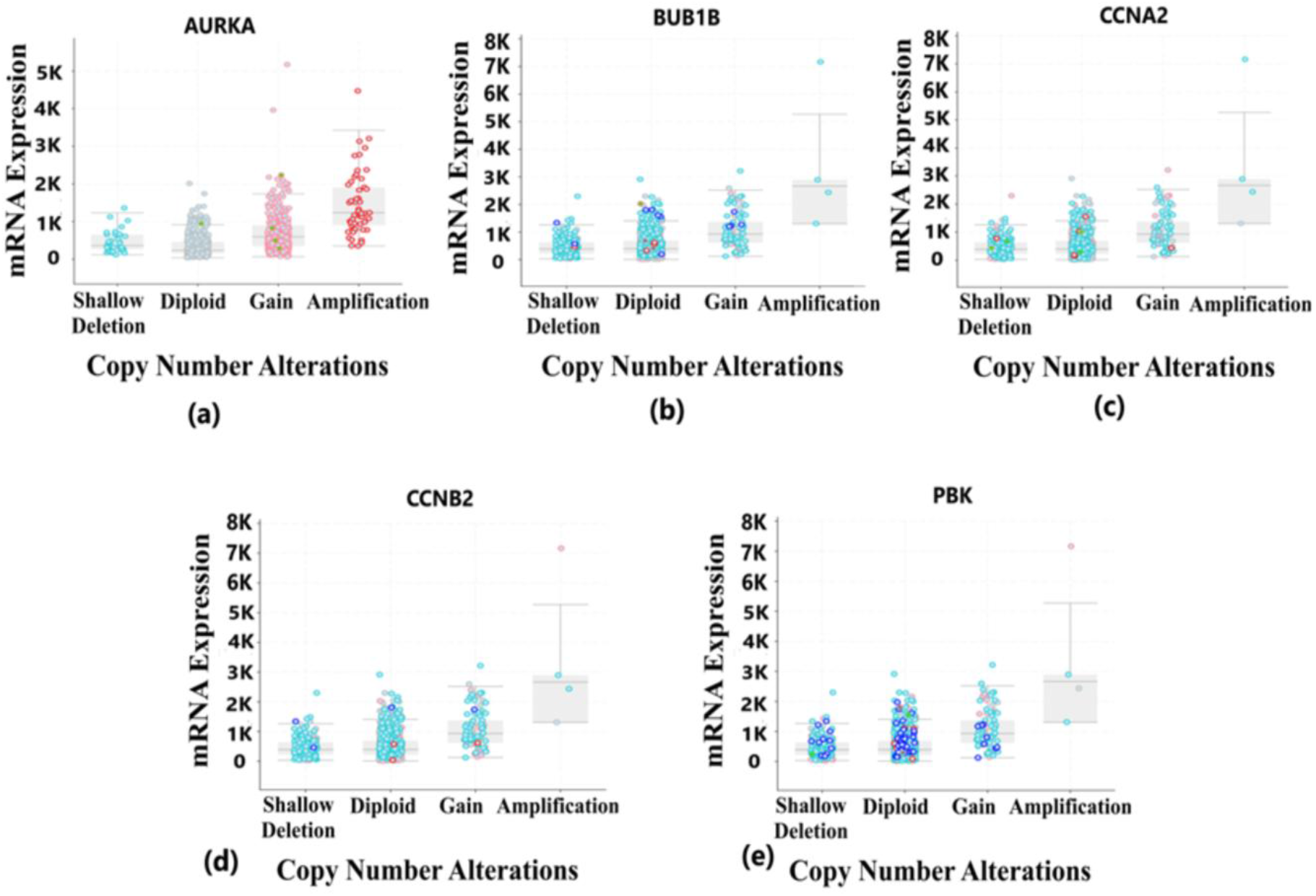
Copy number alterations deals with the deletions and amplification of genetic material fragments. This phenomenon is common in different cancer types and plays a vital role in the development and progression of cancer. This figure belongs to copy number alterations in hub genes of BRCA. (a) Copy number alterations in AURKA gene having most of the changes due to gain and amplification in the genetic materials (b) Copy number alterations in BUB1B gene having most of the changes due to diploid, gain and shallow deletion in the genetic materials (c) Copy number alterations in CCNA2 gene having most of the changes due to shallow deletion, gain and diploid in the genetic materials (d) Copy number alterations in CCNB2 gene having most of the changes due to gain and amplification in the genetic materials (e) Copy number alterations in PBK gene having most of the changes due to gain and amplification in the genetic materials.

### Survival Analysis validation of Prognostic Biomarkers

The aberrant expression of AURKA, BUB1B, CCNA2, CCNB2, and PBK resulted in poorer survival rate of breast cancer patients in the high-risk group having survival rate less than 2 years. The median survival rate was less than 2 years for all the 5 hub genes (Figure 10). For each patient, the risk score was calculated and ranking was done accordingly in the TCGA datasets. Patients were then divided into a high-risk group and a low-risk group. The hazard ratio of the hub genes indicates the risk associated with the survival of the patients (Table 6). The survival rate of the patients was found to be the least in case of overexpressed BUB1B having survival probability of low-risk patients only 48% and those in the high-risk group only 18% and hazard ratio was also highest as compared to other hub genes.

**Table 6.** Table showing the survival analysis results of hub genes in breast cancer.

| S. No. | Gene | Hazard Ratio (CI) |
| --- | --- | --- |
| 1 | AURKA | 1.32 (0.91– 2.42) |
| 2 | BUB1B | 1.85 (1.32 – 2.92) |
| 3 | CCNA2 | 0.49 (0.40 – 0.95) |
| 4 | CCNB2 | 0.62 (0.41 – 1.01) |
| 5 | PBK | 1.26 (0.90 – 2.13) |

**Figure 10.**
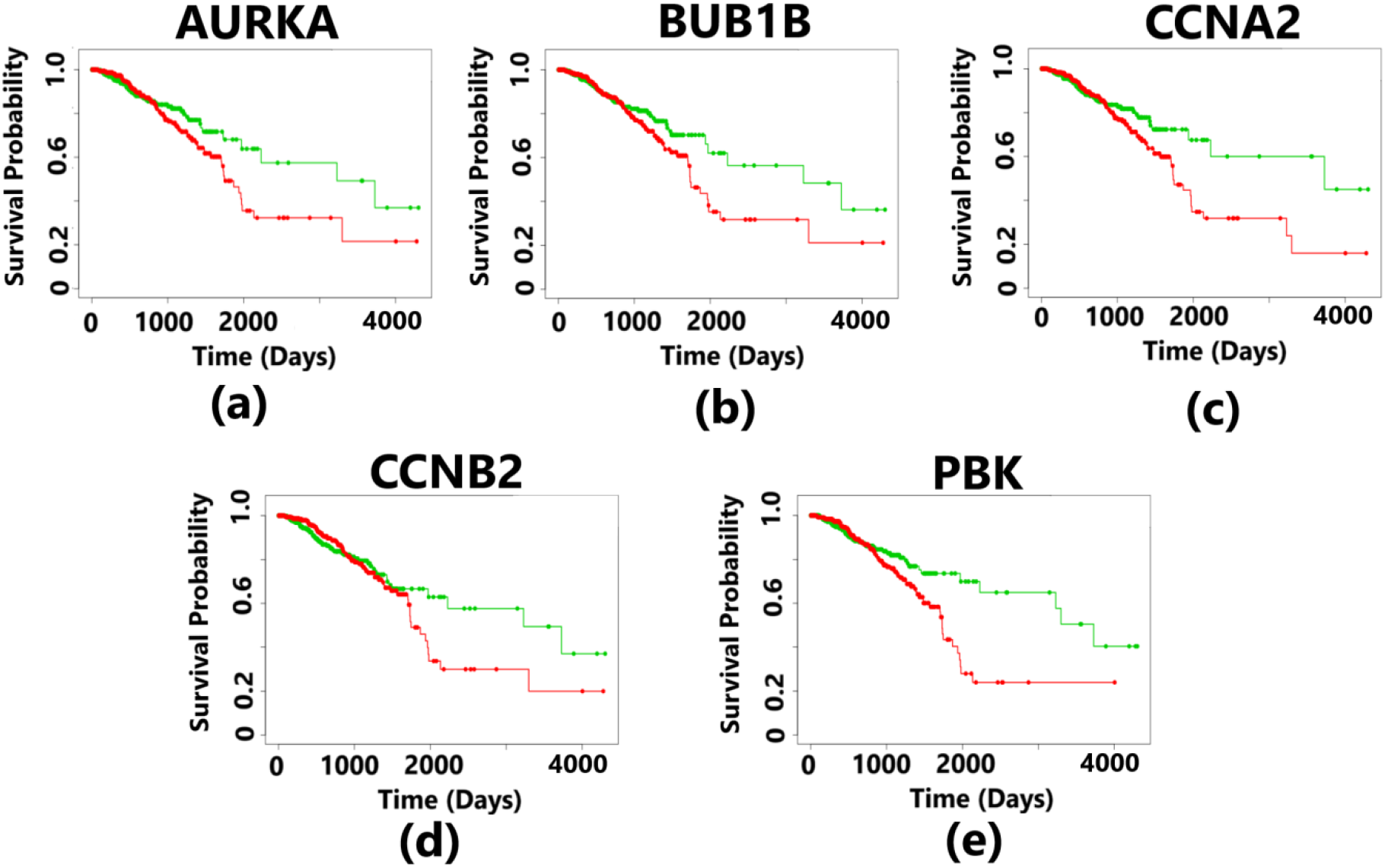
Kaplan-Meier plots showing the survival analysis corresponding to 5 hub genes in breast cancer. The patients were divided into high- and low-risk groups. The overexpression of all the hub genes resulted in the poor survival outcomes which is less than 3 years for the patients suffering from breast cancer. The plot in green color indicates the survival of patients in the low-risk group and the plot in red color represents the survival of patients in the high-risk group. (a) The overexpression of AURKA gene indicate survival probability of 50% for patients in the low-risk group and 25% survival of the patients in the high-risk group (b) The overexpression of BUB1B gene indicate survival probability of 48% for patients in the low-risk group and 18% survival of the patients in the high-risk group (c) The overexpression of CCNA2 gene indicate survival probability of 56% for patients in the low-risk group and 18% survival of the patients in the high-risk group (d) The overexpression of CCNB2 gene indicate survival probability of 54% for patients in the low-risk group and 22% survival of the patients in the high-risk group (e) The overexpression of PBK gene indicate survival probability of 52% for patients in the low-risk group and 23% survival of the patients in the high-risk group

## Discussion

Cancer is a dreadful disease and it costs millions of lives every year, more specifically, breast cancer, which is common among women across the globe. Proper awareness of the biological insight and better understanding of this cancer type through complex networks and signaling might help in early diagnosis and treatment of breast cancer (Vishnubalaji et al., 2019). This in-depth understanding was studies in this research work through transcriptome analysis. The transcriptome analysis paved the way to identify the overexpressed differentially expressed genes that could be potential prognostic biomarkers of breast cancer that could help in prohibiting the tumorigenesis and metastasis of breast cancer. According to the previous studies, the identification of cancer patients with high risk of breast cancer is important to provide effective and specific treatment. These above discussed gene expression profiling concepts will aid in the identification of novel prognostic biomarkers with greater accuracy (Lee et al., 2018). The identified biomarkers could regulate the study of survival analysis of the patients using KM plot based on which the survival probability could be predicted thereby proving these biomarkers as potential therapeutic involved the identification of differentially expressed genes (DEGs) through transcriptomic approach and subsequently obtained the protein-protein interaction network by utilizing these DEGs to identify the most prominent hub genes (prognostic biomarkers) viz. AURKA, BUB1B, CCNA2, CCNB2, and PBK, and these hub genes obtained were found upregulated (based on log_2_fold change value) in breast cancer. Pathway enrichment analysis further showed the biological processes and pathways in which these biomarkers were enriched. The survival analysis predicted poorer prognosis of the patients suffering from these cancer types due to the overexpression of these prognostic biomarkers. The promoter methylation analysis further showed these biomarkers to be hypomethylated breast cancer and that could be a probable cause of spread of breast cancer and development (Hoffman et al., 2005). Moreover, the analysis of genetic alterations that provides information pertaining to variations in prognostic biomarkers could furnish how these changes aid in the progression and metastasis of cancer and its detection, diagnosis and prognosis (Herceg et al., 2007). This genetic alterations in the form of mutations and copy number alterations provided in-depth understanding of genetic changes in the biomarkers that resulted in the tumorigenesis and metastasis of breast cancer in patients.

The five potential prognostic biomarkers i.e. Aurora Kinase A (AURKA), BUB1 Mitotic checkpoint serine/threonine kinase B (BUB1B), Cyclin A2 (CCNA2), Cyclin B2 (CCNB2), and PDZ binding kinase (PBK) were upregulated in breast cancer. These genes were enriched in some of the important biological processes that include mitotic cell cycle, cell division, regulation of mitotic cell cycle, and chromosome segregation. Chromosome segregation is particularly important biological process due to its relation in the development and progression of cancer. The errors introduced in chromosome segregation during mitosis leads to chromosomal instability, which is responsible for tumorigenesis, cancer metastasis and poor prognosis in cancer patients (Bakhoum et al., 2018). The abnormal count of chromosomes due to genomic instabilities plays a pivotal role in tumorigenesis and cancer metastasis (Thompson et al., 2013). The important KEGG pathways that participated in tumorigenesis and metastasis showed the enrichment of the biomarkers in p53 signaling pathway, cell cycle, oocyte meiosis, progesterone-mediated oocyte maturation, glucagon signaling pathway, and PPAR signaling pathway. In this study it was observed that the potential biomarkers are overrepresented in cell cycle KEGG pathway. This improper regulation of cell cycle may result in uncontrolled cell multiplication and this phenomenon leads to tumorigenesis and cancer metastasis (Novitasari et al., 2021). The two other important KEGG pathways in which the biomarkers were enriched are oocyte meiosis and progesterone-mediated oocyte maturation. In the meiosis process, two more rounds of chromosome segregation (Meiosis I and Meiosis II) are followed by a single round of DNA replication (Mehlmann et al., 2012). At G2 of meiosis I, oocytes are naturally arrested and this arrest is broken by the encounter to the progesterone which is a steroid hormone. This persuades the maturation of the oocyte and the two meiotic division cycles process to be resumed (Mahrous et al., 2012; Shao et al., 2013). So, it may be inferred that the cell cycle process might get affected due to abnormal regulation of meiosis and oocyte maturation. Moreover, this change in cell cycle had a negative impact on normal activities in the human body resulting in increased risks of suffering from different cancer.

AURKA, which belongs to serine/threonine kinase family, is responsible for activating the process of cell division through mitosis regulation. It is an oncogene and promotes tumorigenesis and metastasis in different cancer types and it is due to this property that AURKA is treated as potential target in cancer treatment (Du et al., 2021). Its overexpression in cancer cells as compared to normal cells resulted in probable development and spread of breast cancer. This gene is related to cell cycle progression and due to its overexpression and oncogenic role and hence its inhibition might lead to regression of breast cancer (Siggelkow et al., 2012). As compared to the normal samples, the tumor samples had higher methylation and the gene was hypomethylated (beta value - 0.034) and this hypomethylation caused genetic instability which is the primary reason for development and metastasis of breast cancer. Further, the genetic alterations analysis provided information about the different mutations that happened in AURKA and caused its upregulation. These mutations were of missense type and occurred at 5 different mutation sites (S98N, S4Y, S89C, A81V, and L26V) and also this gene has been identified in amplified regions due to gene amplification. This combined effect of mutation and amplification caused genetic alterations and in turn resulted in altered phosphorylation. This alteration in phosphorylation post-transcriptional modifications (PTMs) affected many significant pathways in which AURKA gene was enriched such as cell cycle and played a key role in breast cancer growth and metastasis. So, these altered phosphorylation that are strongly associated with breast cancer could be a potential target for development of suitable anti-cancer drugs that can inhibit the progression and metastasis of breast cancer. The diploid and gain copy number alterations found in this gene also played a role in the development and progression of breast cancer. The survival analysis showed poor prognosis in case of AURKA having hazard ratio > 1 (1.32). The patients in the low-risk group have higher survival probability (50%) than those in the high-risk groups with survival probability 25%. The overexpression and poor survival rate indicate that this gene to be a potential predictive biomarker for early detection and diagnosis of metastatic breast cancer.

BUB1B gene belongs to the serine/threonine kinase family. Its genomic location is 15q15.1. This gene plays a vital role in encoding a kinase which is involved in spindle checkpoint function. Due to its localization to the kinetochore, it serves an important purpose of inhibiting the anaphase-promoting complex/cyclosome (APC/C). This process delays the onset of anaphase and ensures proper chromosome segregation. The impairment in the spindle checkpoint function has resulted in many cancer forms. In this study, BUB1B was overexpressed in breast cancer which showed the probability of breast cancer metastasis. In breast cancer metastasis, the chromosomal instability was found to be the main cause and this defect pertains to imperfection in mitotic spindle checkpoints. This process is related to the overexpression of BUB1B gene (Yuan et al., 2006). Another study reported that overexpression of BUB1B gene caused a decrease in the survival probability of the patients suffering from breast cancer and it also resulted in its metastasis (Koyuncu et al., 2020).

The analysis of DNA methylation showed that the promoter methylation level of tumor samples was lower as compared to normal samples indicating higher expression of this gene in case of breast cancer (beta value – 0.125). The genetic alterations in this gene was caused due to missense mutation at 2 sites (Q460E and L669P) and nonsense mutation at a single site (S564*). Another genetic alteration that affected this gene was amplification where this gene was identified in the amplified regions, although the frequency of amplification was very less (0.32%) in the breast cancer patients affected due to the overexpression of this gene in contrast to mutation (0.39%) and deep deletion (1.12%). Deep deletion, and particularly FS deletion at D989Mfs*13 also altered the normal functioning of this gene. These three genetic alteration types resulted in the genomic instability and this in turn resulted in the tumorigenesis and metastasis of breast cancer. The copy number alterations in this case were gain and diploid CNA as these two are most prominent in producing cancer. The survival analysis of BUB1B pertaining to breast cancer showed survival probability of 48% in case of low-risk group patients and 18% in case of high-risk group patients. The hazard ratio was 1.85 which was very high and proved the overexpression and poor prognosis of this gene to be a potential prognostic biomarker for breast cancer.

CCNA2 is a protein coding gene which belongs to the cyclin family. This family is considered as highly conserved and its members are mainly involved in regulating the cell cycle. The cyclin-dependent kinase 2 was activated through the binding of this protein and thus helps in promoting transition through G1/S and G2/M. It plays a prominent role in progression and distant metastasis of breast cancer and in turn as a biomarker (Gao et al., 2014).

The findings of the following study shows that CCNA2 was overexpressed in case of breast cancer and since this gene has an oncogenic role in cancer (Gan et al., 2018), it participated in the tumorigenesis and metastasis of breast cancer. The promoter methylation level of tumor samples is higher than normal samples with beta value 0.10. This value indicates that the CCNA2 gene was hypomethylated in breast cancer leading to the speedy tumor progression and metastasis. The genetic alterations that are involved in the overexpression of CCNA2 include mutation and amplification. The missense mutation at 4 mutation sites (R112C, L315P, M189I, and V85F) was primarily responsible for changes produced in the expression level of this gene. The other genetic alteration i.e. amplification was related to increased growth of breast cancer cells and further assisted in its metastasis due to the upregulation of CCNA2 gene. The copy number alterations that were associated with this gene include diploid and shallow deletion CNA and both of these are already discussed to promote tumor growth and metastasis. The survival analysis demonstrated that the overall survival probability of the patients in the low-risk group was 56% for 1 year and that in the high-risk group it was only 18%. The hazard ratio was < 1 (0.49) and this showed that the overexpression of this gene was comparatively less effective in case of breast cancer as compared to other biomarkers. However, the survival probability, particularly in case of high-risk group was associated with poor prognosis and hence this gene could be a significant predictive biomarker for the diagnosis and inhibition of breast cancer tumorigenesis and metastasis.

CCNB2 gene belongs to B-type cyclin family. This gene has been found to play a vital role in G2/M transition in eukaryotes through the activation of CDC2 kinase (Shubbar et al., 2013). The upregulated CCNB2 gene was found to be responsible for developing Lymphovascular Invasion (LVI), a key step in breast cancer metastasis (Alzohani et al., 2022).

The overexpression and oncogenic role of CCNB2 gene was responsible for the metastasis of breast cancer. Studies related to breast cancer suggested that CCNB2 was abnormally expressed in this cancer type and particularly in case of triple negative breast cancer. This abnormal behavior resulted in tumorigenesis and metastasis of breast cancer proved its potential as a suitable therapeutic target for the treatment of triple negative breast cancer (TNBC) (Wu et al., 2021). This study also demonstrated the overexpression of CCNB2 and this had an adverse effect on the normal functioning of the cells and hence the breast cancer cells metastasized. Moreover, the promoter methylation level showed that the expression level of tumor samples was lower as compared to the normal cells thereby indicating higher expression level of this gene in case of metastatic breast cancer (beta value – 0.06). The genetic alterations consisted of amplification and deep deletion. These two alterations took part in the promotion of tumor growth and metastasis. The survival analysis obtained from the multivariate cox analysis indicated hazard ratio < 1 (0.62). This value shows that the expression level of this gene was in a controlled manner. The survival probability was 54% in low-risk group of patients and those in the high-risk group has survival probability of 22%. This result of survival was well acquainted with the hazard ratio that was obtained for this gene. Hence, this gene can significantly assist in predicting the poor survival of the patients and could act as a promising therapeutic target for in HCC (beta value – 0.05). The genetic alteration analysis showed that only gene amplification participated in producing the genomic instability of this gene. The survival analysis provided information about the hazard ratio that was greater than 1 (1.27) and this shows the higher expression of CCNB2 as was also shown through promoter methylation analysis. The survival probability in the low-risk group patients was 20% in contrast to that in the high-risk group patients where the survival probability was 42%. This higher expression of CCNB2 as shown by the results of promoter methylation and poor prognosis obtained from the survival analysis demonstrated the efficacy of this gene to be a suitable candidate for prediction, diagnosis and treatment of HCC.

PBK gene belongs to mitogen-activated protein kinase kinase (MAPKK) family. It encodes a serine/threonine protein kinase that is related to the dual specific MAPKK. Overrepresentation of this gene has been involved in the process of tumorigenesis. The overexpression of PBK gene was found to have an association with poor survival of patients in different cancer and this made PBK a suitable prognostic biomarker and a potential therapeutic target (Xu et al., 2019).

In case of breast cancer, the PBK gene was found to be overexpressed and this resulted in the progression and probable metastasis of breast cancer to form GBM and HCC. In one of the latest studies it was reported that the overrepresentation of PBK gene resulted in poor prognosis of patients suffering from breast cancer (Qiao et al., 2022). From the following study, it was further observed that promoter methylation level of tumor samples was higher as compared to the normal samples and this indicated lower expression of this gene in case of breast cancer (beta value – 0.25). The genetic alteration study further demonstrated the involvement of amplification, mutation, and deep deletion in producing the overexpression of PBK gene. The missense mutation at two mutation sites (E203K and F40L) and nonsense mutation at a single mutation site (E295*) showed the genomic instability that caused for the growth and metastasis of breast cancer. Besides mutation, the other two alterations that were responsible for overexpression include amplification and deep deletion (FS deletion at K18Efs*50). The phosphorylation post-translational modification was also altered resulting in further progression of cancer. The survival analysis showed that the hazard ratio was 1.26. The survival probability of patients in the low-risk group was 52% while that in the high-risk group was 23%. The poor prognosis of this gene qualified it to be a suitable indicator for the prediction and diagnosis of breast cancer metastasis.

## Conclusion

The present study provided five potential prognostic biomarkers viz. AURKA, BUB1B, CCNA2, CCNB2, and PBK through the integrated approach of transcriptome and pathway enrichment analysis. This will aid in early diagnosis and treatment of breast cancer and could probably improve the survival analysis of the patients. By proper designing of potential inhibitors for these biomarkers will help immensely in suppressing tumorigenesis and metastasis of breast cancer.

## Notes

### Competing Interest Statement

The authors have declared no competing interest.

